# Influence of anthropogenic light on puma road crossing movement

**DOI:** 10.1101/2022.10.28.514303

**Authors:** Eric S. Abelson, Jeff A. Sikich, Seth P.D. Riley, Daniel T. Blumstein

## Abstract

Wildlife may be attracted or repelled by anthropogenic stimuli – by understanding these responses we gain powerful tools to modify their behavior. Anthropogenic lighting is ubiquitous but little is known about how it influences carnivores. We focused on pumas (*Puma concolor*) living around Los Angeles, California for which vehicular collision is an important source of mortality. We used GPS collars to track their movement around roads and combined this with ground-level and remotely sensed data of human generated nighttime light to see how that influenced activity around roads. There are multiple landscape level scales, and metrics of light relevant at each scale, that influence the perceptual landscape of wildlife and thus these different scales must be considered. We found that, at a broad spatial scale, pumas crossed roads more frequently in areas with low levels of anthropogenic light (controlling for distance to urbanization). However, while pumas crossed roads in darker parts of the landscape at a broad-landscape scale, we did not find a statistically significant relationship between puma eye-level light intensity and predicted road crossing locations.

Regarding pumas and nighttime light, it appears that having dark swaths of land, at a broad spatial scale, is important. This is important for puma conservation and road design as little is currently known about wildlife movement response to nighttime light and the ideal placement for mitigation structures. Ultimately, anthropogenic light at night is a landscape aspect that needs to be better understood and integrated into conservation as the human footprint increases.

## 1. Introduction

Wildlife rely on their senses to collect information about their landscape that, in turn, influences how they move through their environment and use resources (Smith et al. 2015). While behavioral responses to environmental cues have been honed over evolutionary time, there are many stimuli (especially in landscapes modified by humans) that have become decoupled from their historical meaning (Robertson & Hutto 2006). Directly studying wildlife perceptual abilities is difficult, but we can infer perception and downstream behaviors from studying wildlife movement and habitat selection. Many species live in highly fragmented habitats and roads crisscross their home ranges. Because roads are a significant source of mortality for some species, understanding how anthropogenic features influence road crossing behavior should have important conservation implications.

We studied how anthropogenic lighting influenced movement of pumas (*Puma concolor*) across roads in the mountains around Los Angeles, California, USA. Roads are among the leading causes of puma mortality in Southern California and in the Santa Monica Mountains (Vickers et al. 2015; Benson et al. 2020). Nighttime light dominates the visual perceptual landscape and impacts nocturnal wildlife species’ behavior and movements (Zollner & Lima 1999; Rich & Longcore 2013). Anthropogenic contributions to nighttime light are particularly disruptive and are known to influence wildlife in many ways including causing disorientation, shifts in activity patterns, and disrupted navigation (Rich & Longcore 2013). By identifying how light affects road crossings, we gain tools that can be used to both attract and repel (Greggor et al. 2020) pumas (e.g., make busy roads appear unsafe and make dedicated wildlife crossing structures appear safe).

Our study area of Southern California’s Santa Monica Mountains is well-suited to examining how puma movement around roads is influenced by anthropogenic light; it is a natural landscape with varying levels of anthropogenic encroachment and is fragmented by roads. To assess the role that anthropogenic light plays on modifying puma road-crossing, we explored the relationship between light and puma road-crossing behavior at two spatial scales using three distinct metrics; we hypothesized that the number of times pumas crossed roads in particular locations, or road crossing intensity, will be influenced by nighttime light intensity. Using a puma GPS-collar dataset for 17 pumas collected between 2002 and 2013, we examined road crossing intensity and light measurements from both puma eye-level at the local scale, as well as at the broad landscape scale using remotely sensed light levels. Puma-eye level measurements are split into a “horizon-view” perspective that captures light above the road verge (i.e., what lies across the road) and a “road-view” perspective that examines the amount of light on the road surface itself.

## 2. Materials and Methods

All statistical analyses were completed in program R 4.0.2 (R Core Team 2020).

### 2.1 Study area

The study area lies within the Mediterranean Ecosystem of the Santa Monica Mountains National Recreation Area in Los Angeles and Ventura Counties in California. Dominant vegetation communities included chaparral, coastal sage scrub, oak woodlands, and introduced grasslands. Human land uses included commercial and industrial areas and residential areas from high (greater than five houses per two acres) to low (fewer than one house per two acres) density, although within the core Santa Monica Mountains themselves, housing is generally low-density. Overall, the Santa Monica Mountains are isolated from other natural areas: specifically, puma habitat is hemmed in to the east and north by urbanization and freeways, to the west by agricultural areas in the Oxnard Plain, and to the south by the Pacific Ocean. To avoid the influence of dense urbanization, the study focused on the core of the Santa Monica Mountains, consisting of extensive wildlands with interspersed housing (Fig. 1).

**Figure 1.**
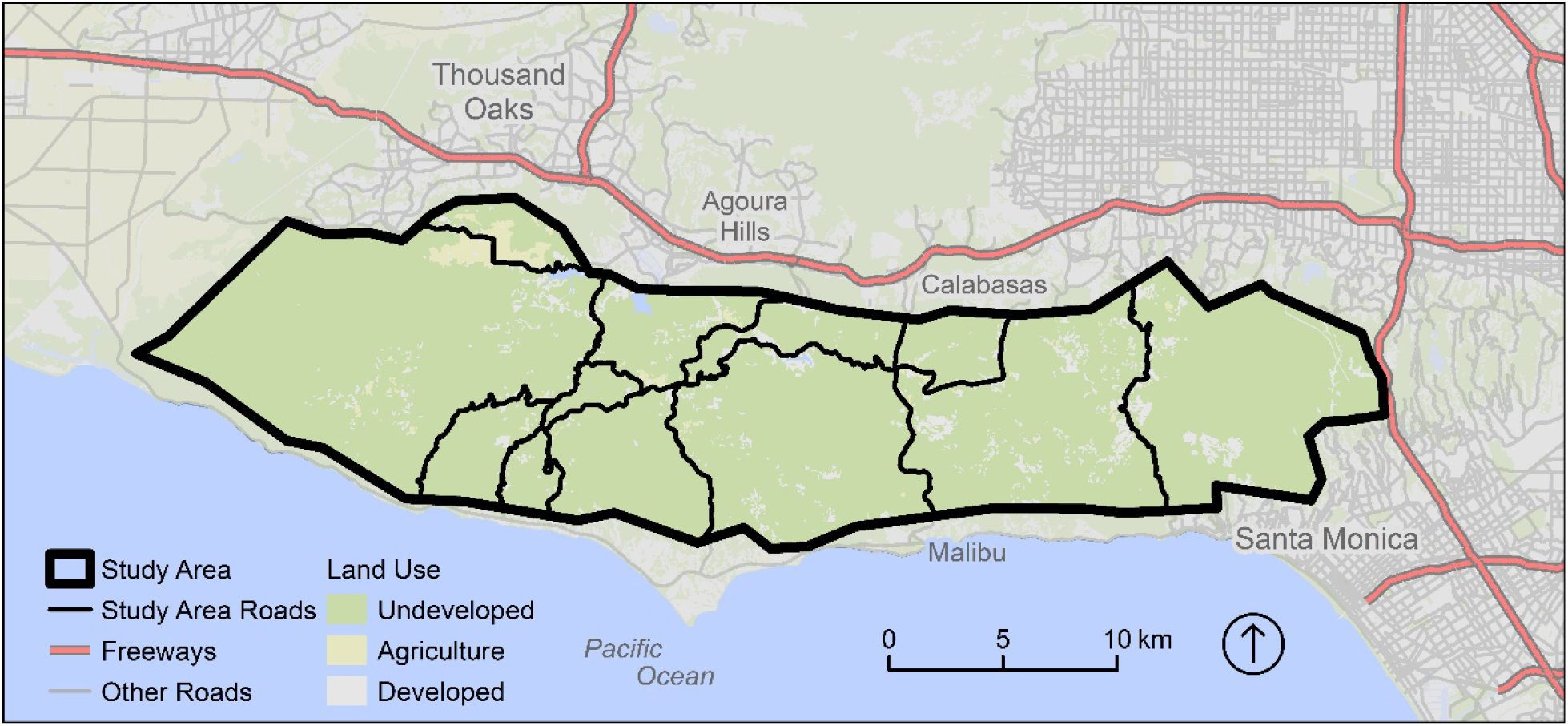
Study area. Puma study area and roads used in broad landscape-scale light analyses. This study area is situated in Southern California’s Santa Monica Mountains National Recreation Area.

Using the StreetMap07 roads layer, (GDT Dynamap 2014) we identified 109 km of secondary road in our study area (U.S. Census Bureau’s Census Feature Class Code, hereafter FCC, classes A30-A38). Landscape level analyses described below were performed by dividing the 109 km of road into 68 1.6 km segments; only road segments that were 1.6 km long with the same FCC class and road name were included in analyses.

### 2.2 Light measurements

We performed four analyses to explore the nighttime light environment of pumas – two at a broad landscape scale (i.e., ~400 square meters), and two from puma eye-level at the road verge.

To examine anthropogenically-created nighttime light (radiance, W cm^-2^ sr^-1^) at a broad spatial-scale we used moonless and cloudless composites collected by the Visible Infrared Imaging Radiometer Suite Day-Night Band (VIIRS) that was aboard the NASA/NOAA Suomi-NPP satellite (Elvidge et al. 2017); this metric of nighttime light has been used in other wildlife studies (Hu et al. 2018; Levin et al. 2020). In our study area each VIIRS pixel measured approximately 380 m by 470 m. For each 1.6 km segment of road, we extracted and calculated the mean VIIRS light levels at 161 m increments.

Light at puma eye-level was measured using methods developed by light engineer Peter Hiscocks (Hiscocks 2011, 2013a, 2013b, 2014) where RAW format image data were calibrated using a professional spectroradiometer and light integrating sphere and then converted to luminance (candela/m^2^). Canon D30 cameras (Canon, Melville, NY), using the Canon Hack Development Kit (CHDK; Russell 2016) on camera software (Russell 2016), photographed the scene across roads from a puma-eye perspective (approximately one meter above ground-level) at 74 field sites throughout the night. At each field site a nighttime photo was selected that was not obscured (by rain or vegetation), had no light from passing vehicles, was taken during astronomical twilight (i.e., center of the sun was geometrically 18 degrees below the horizon), and the moon was at least one hour from rising or one hour after it had set. We also extracted VIIRS light (log_10_ radiance) at each puma eye-level field site and calculated minimum, maximum, mean, and standard deviation. While VIIRS light levels (log_10_ radiance) cannot directly be compared against puma eye-level light metrics (log_10_ luminance), summary statistics can be used to identify where puma eye-level sites fall in the general range of VIIRS light levels from the broad-landscape scale analyses.

We attached light measuring equipment to power-poles owned by Southern California Edison. Sites were established without knowledge of predicted puma crossing intensity so as not to bias measurements. Equipment was placed (no closer than 200 m to each other) along the following roads in the Santa Monica Mountains study area (number of sites per road follow parenthetically): Mulholland Hwy. (36), Las Virgenes Rd./Malibu Canyon Rd. (13), Old Topanga Canyon Rd. (9), Topanga Canyon Blvd. (9), Decker Rd. (5), Colina Dr. (1), Stokes Canyon Rd. (1). Coordinates, light measurements, and site data are available in the supplemental information (Appendix S1).

Resulting lightscape data were split into two components: 1) horizon-view: mean luminance for pixels falling above the distant road verge to quantify light in the view-shed of an approaching puma and 2) road-view: pixels below the distant road verge mean luminance; a proxy for how well an approaching puma could see the road itself and to discern nearby objects. CHDK software was installed on each camera to control the camera and to generate RAW format images (Hiscocks 2014). Resulting RAW images were processed in ImageJ (Rasband n.d.) split into the above- and below-distant road verge components. Using the workflow and formulas described by Hiscocks (Hiscocks 2013b, 2014) luminance was calculated from pixel values. Calibration coefficients were determined by using a research-grade light integrating sphere and a PR-650 spectroradiometer (Photo Research, Chatsworth, CA) in a controlled lab setting (Appendix S1).

### 2.3 Modeled road crossing locations

A puma movement dataset was constructed using GPS locations from seventeen pumas, using GPS collars, tracked between 2002 and 2013; this dataset consisted of 53,825 fix locations. To identify the most probable locations where pumas crossed roads we modeled least cost movement paths (Wade et al. 2015) starting from each puma coordinate to the subsequent coordinate following the methods in Lazo-Cancino et al. (Lazo-Cancino et al. 2017). We used least cost paths because simple straight-line paths between fix locations were biologically unrealistic, did not follow topography or vegetation that was preferentially used by pumas, and may result in an unlikely number of road crossings along curvy roads. To minimize uncertainty, any paths generated from puma location pairs that were separated by more than 120 minutes were excluded from the analysis. Movement paths and roads were then examined to find intersections: we found 2,459 locations where least cost paths intersected roads in the study area from fifteen pumas (two of the seventeen pumas were not predicted to have crossed roads in the study area) and these coordinates were used as predicted puma crossing locations in the following analyses.

Following Zeller et al. (2014), a resistance surface was parameterized with land cover, which included both vegetation types (National Park Service 2014) and human land use types (SCAG 2005), slope data (USGS 2015), and puma location data. We calculated pixel resistance values in our study area based on probability of use calculated by examining empirically “used” habitat (determined by using a 30m buffer around each puma fix location) and “available” habitat using Pareto distributions and kernels (Zeller et al. 2014).

### 2.4 Proximity to urban areas

Light levels are often, but not always, correlated with urbanization. We hypothesized that predicted road crossing intensity will vary with proximity to urbanization. We included distance to urbanization in our models and tested for collinearity between predictor variables to ensure that proximity to urbanization and light levels were not highly correlated. The landscape-scale analysis used distance to urbanization calculated at 161 m intervals along each road; for each 1.6 km road-segment we calculated the mean distance (in meters) to the nearest urban area (defined as commercial, industrial, or residential with greater than five houses per two acres) across 161 m points that fell within that road segment. For puma eye-level analyses we used the distance from each field site to the nearest urban area. We used the gDistance function (*rgeos package* 2020) to calculate the distance between each point of interest (i.e., either field site location or 161 m point) and the closest urban polygon from (“Santa Monica Mountain Recreation Area vegetation map, super-generalized version.” 2015).

### 2.5 Broad spatial-scale response to light analysis

We examined if predicted puma road crossing locations are biased by an interaction between light conditions and road type while controlling for the effects of proximity to urbanization; we hypothesized that predicted road crossing intensity would vary with light intensity. To examine puma response to broad-landscape level light we examined number of predicted crossings per 1.6 km segment of road (hereafter crossing intensity).

To examine the relationship between puma road crossing intensity and light levels we fitted a linear model using the lm function after log_10_ transforming both predicted road crossing quantity and light levels. Crossing intensity was the dependent variable while independent variables were light level (VIIRS), and proximity to urbanization. To check for collinearity between independent variables we used a generalized variance-inflation factor analysis using the “vif” function in the car package (*car package* 2020) with crossing intensity as the dependent variable and proximity to urbanization and light level (VIIRS) as independent variables. We used the gvlma function (*gvlma package* 2019) to ensure the model met model assumptions for skewness, kurtosis, and heteroscedasticity.

### 2.6 Assessing puma baseline light level exposure vs. light levels at predicted road crossing locations

To examine if light levels at predicted road crossing locations were directly associated with baseline levels of light that pumas were exposed to, we compared the two light distributions to each other. Baseline light exposure was calculated as light levels (extracted from VIIRS satellite data) at each of 53,825 puma GPS collar locations. Light levels from the 2,459 modeled road crossing locations, also extracted from VIIRS, formed the distribution of modeled road crossing light levels. We used a two-sample Kolmogorov-Smirnov test (Conover 1998) using the ks.test function where the null hypothesis is that light levels at modeled road crossing locations and baseline light levels, locations where pumas are not crossing roads, are pulled from the same distribution. If pumas were crossing roads in the same parts of the landscape that they moved more generally, both baseline and predicted road-crossing light levels would follow the same light intensity distribution, and we would accept the null hypothesis. Rejecting the null hypothesis suggests that light at predicted road crossings is not simply an artifact of where pumas happen to be on the landscape in that light levels where pumas cross roads is different than where they generally move.

### 2.7 Puma eye-level response to light analysis

We examined crossing intensity and light intensity on the road (road-view) and above the road surface (horizon-view). For these analyses, we calculated crossing intensity as the number of modeled puma crossings within 50 m of each site (i.e., along a 100 m stretch of road with the site being in the middle; table 1). Road-view and horizon-view light intensity values were calculated as described in the light measurements section.

**Table 1.**
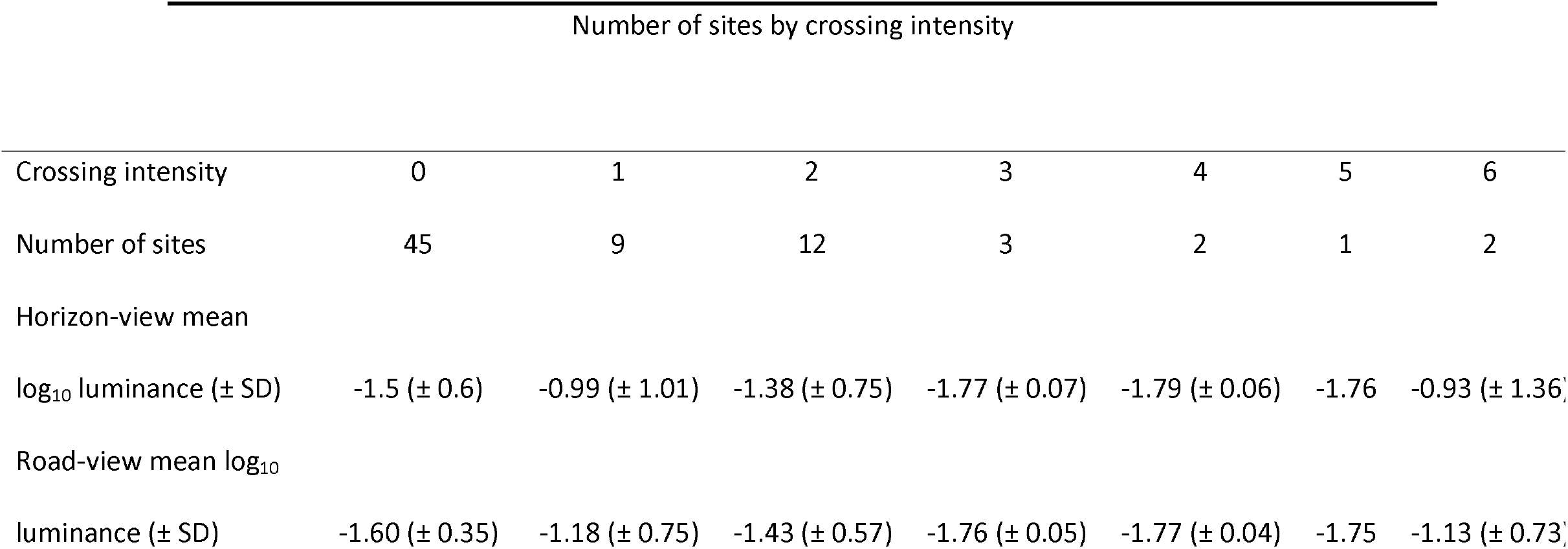
Number of field sites, for which eye-level light intensity was measured, at each level of predicted crossing intensity. Field sites are locations along roads in California’s Santa Monica Mountains National Recreation Area where light was measured across the field-of-view from a puma perspective. Light levels were measured, and the mean luminance across the field-of-view is reported here for both the area above the road verge (horizon-view) and below the road verge (road-view). Crossing intensity was calculated as the number of predicted puma road crossings within the 100 m segment of road around the field location (i.e., 50 m on either side of the field site).

To examine the relationship between puma road crossing-intensity and horizon-view and road-view light intensity, we fitted two linear models after log_10_ transforming light intensity and log_10_ transforming road crossing after adding one (because there were sites with zero crossings) so that methods were analogous across road-view, horizon-view, and broad spatial-scale analyses. The dependent variable was crossing intensity and independent variables were light level and proximity to urbanization. No interaction term was used in horizon-view and road-view analyses, unlike in the broad spatial-scale analysis; all but two sites were along large secondary roads resulting in no effective variability in road type.

### 2.8 Capture and monitoring

Pumas were captured using foot cable-restraints (also known as Aldrich foot-snares), baited cage-traps, or by treeing them with trained hounds during 2002-2013. We immobilized pumas with ketamine hydrochloride combined with either xylazine hydrochloride or medetomidine hydrochloride administered intramuscularly. Captured animals were monitored for the duration of the time they were immobilized. Adults and subadults were then fitted with a GPS radio-collar (Followit AB, Simplex and Tellus models, Stockholm, Sweden; North Star Science and Technology LLC, Globalstar Tracker model, King George, Virginia, USA; or Vectronic Aerospace, GPS Plus model, Berlin, Germany) equipped with a VHF beacon. Fix schedules for the GPS radio-collars varied but most of them were programmed to obtain either one or two locations during the day and between five and seven nighttime locations per 24 hours. Capture and handling was conducted under California Department of Fish and Wildlife Permit SC-0005636, and the protocol was approved by the National Park Service’s Institutional Animal Care and Use Committee.

## 3. Results

### 3.1 Broad spatial-scale response to light analysis

We found that puma road crossing-intensity decreased as landscape-level light intensity (VIIRS radiance) increased after accounting for proximity to urbanization [β = −0.47, F(1, 65) = 6.43, p = 0.01] (Fig. 2). We did not find a significant main effect of distance to urbanization [β = 0.0002, F (1, 65) = 1.38, p = 0.24]. This model met model assumptions for skewness, kurtosis, and heteroscedasticity. We also find that independent variables were not collinear (GVIF equal to 1.04 for each variable).

**Figure 2.**
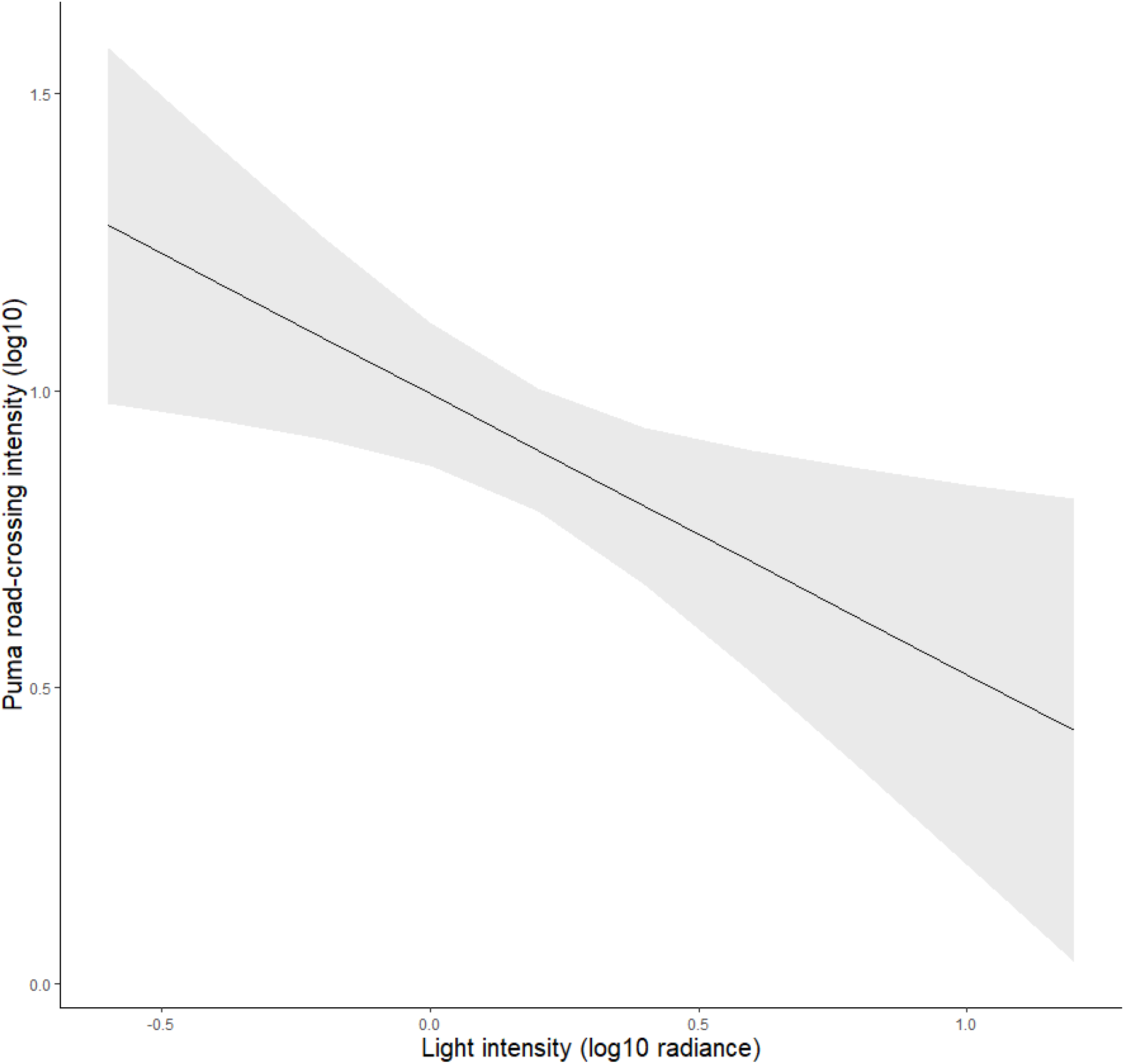
Puma road-crossing intensity as predicted by light intensity and proximity to urbanization. Pumas in the Santa Monica Mountain National Recreation Area between 2002-2013 crossed roads more frequently using segments of road with less human-generated light at night. Model output showing puma road-crossing intensity, per 1.6 km segment of road, decreases as light intensity increases across roads after accounting for proximity to urbanization (p = 0.01; shaded area is 95% CI).

### 3.2 Puma eye-level response to light analysis

We found no relationship between puma road-crossing intensity and puma eye-level horizon-view light intensity [F(1, 71) = 0.81, p = 0.37] or road-view light intensity [F(1, 71) = 1.89, p = 0.17]. We also found no relationship between puma road-crossing intensity and proximity to urbanization either in the horizon-view analysis [F(1, 71) = 0.41, p = 0.52], or the road-view analysis [F(1, 71) = 0.63, p = 0.43] (Fig. 3). For reference, VIIRS light levels (log_10_ radiance) at eye-level sites ranged from −0.27 to 1.12 with a mean of 0.276 (SD 0.29).

**Figure 3.**
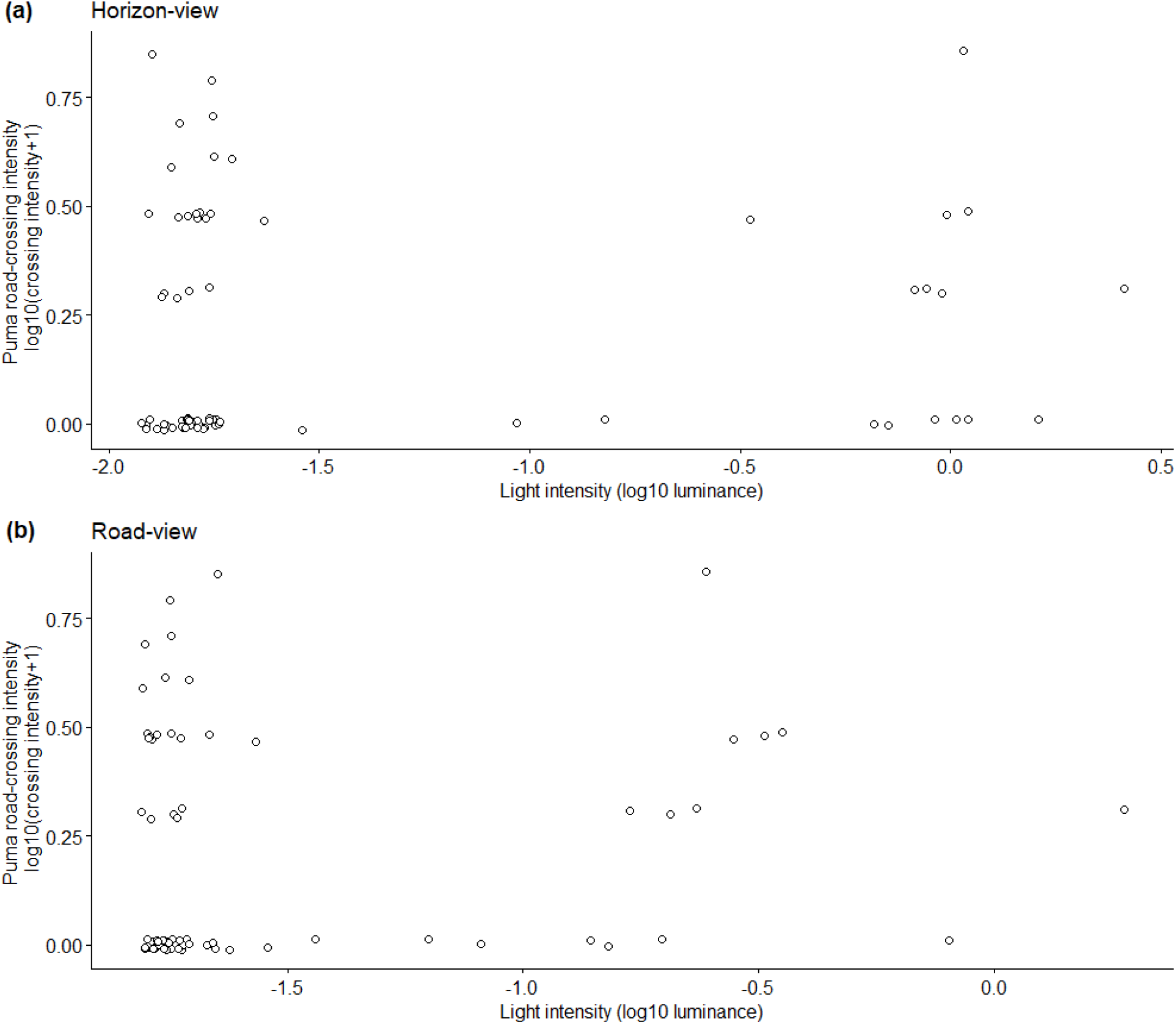
Puma road-crossing intensity as predicted by puma eye-level light intensity. Puma road-crossing intensity (within 50 m of a light-measurement field site) is not significantly correlated with average light levels on the horizon or road surface from a puma’s perspective. To visualize many overlapping points with zero modeled crossings the data points have been jittered (i.e., a small amount of noise has been added to the crossing intensity and light level values so that points fall near but not on top of each other).

### 3.3 Puma baseline light level exposure vs. light levels at modeled road crossing locations

Using a two-sample Kolmogorov-Smirnov test we found evidence to reject the null hypothesis that light levels at modeled puma road crossing locations were from the same distribution as baseline puma locations (D = 0.34, p < 0.0001). As such, light distributions are different when comparing locations where pumas were predicted to have crossed roads against puma non-road crossing locations. Light levels (log_10_ radiance) for all puma GPS locations ranges from −0.63 to 1.90 with a mean of −0.12 (SD = 0.28). Light levels (log_10_ radiance) for modeled crossing locations ranged from −0.53 to 1.10 with a mean of 0.074 (SD = 0.31).

## 4. Discussion

Human-caused landscape alteration is widespread, and disruptions reach the nighttime sky (Rich & Longcore 2013). While there are many examples of ways in which human-caused light at night have deleterious impacts on wildlife, ranging from disorientation to disruption of foraging behavior, there is much that is still unknown, especially in mammals (Rich & Longcore 2013). This lack of knowledge can be especially problematic for conservation biology which focuses on preserving biodiversity in the face of challenges presented in anthropogenic landscapes (Soulé 1985). Road ecology, aspects of which target avenues for protecting both humans and wildlife, is one such area that could benefit from better knowledge of wildlife response to nighttime lighting.

Roads isolate, hamper movement of, and limit gene flow in wildlife (Riley et al. 2003, 2006, 2014b), and road surface area is increasing. Mitigation structures, safe passages for wildlife under or over a road (Ng et al. 2004), can reduce some of the damage that accompany roads – to people, wildlife populations, and the ecosystems in which wildlife play a vital role (Forman et al. 2003; Riley et al. 2014a). While mitigation structures have and continue to be erected the world over, this manuscript addresses a current gap in knowledge: beyond the availability of safe passage from one side of the road to the other, what is the best location to situate these structures and what can be done to alter conditions near the entrance of mitigation structures to encourage use.

Anthropogenic light at night is known to influence wildlife movement and navigation (Rich & Longcore 2013). For example, nighttime lights are known to lead to mass casualties in birds (Rich & Longcore 2013) and sea turtles, emerging from eggs on the beach, move toward city lights and certain death instead of toward the ocean (Witherington 1997). Additionally, eye physiology in nocturnal animals may lead to night blindness in the face of some human generated nighttime light (Rich & Longcore 2013). Previous research has examined the role of nighttime lighting, roads, and wildlife, but this past work focuses generally on either the ability of vehicle drivers to see wildlife (with the possibility of then avoiding a collision; Reed & Woodard 1981) or temporary blindness in wildlife on or near roads (Rich & Longcore 2013). Little is known regarding wildlife response to light at the roadside, and if lighting conditions alter where wildlife choose to approach, and subsequently cross, roads.

We found that landscape-level human-created light at night was correlated with modeled puma road crossing locations after accounting for proximity to urbanization; specifically, we find increased crossing intensity at darker sections of road. This result was not an artifact of baseline puma movements; light levels at modeled road crossing locations are not simply a byproduct of baseline light levels that pumas experience generally (i.e., we find that the distribution of light intensity at modeled road-crossing locations do not follow the same distribution as light intensity at all GPS puma locations generally).

When examining light at puma eye-level though, we found no statistical relationship between modeled road crossing location and average light intensity. It is important to note that the lack of statistical significance does not necessarily indicate that there is no effect of light on local puma movements.

While these results may initially seem contradictory, we believe that these findings instead 1) underscore the importance of considering light in analyses of puma movement, 2) deepen our understanding of how pumas may respond to light differently at different scales, and 3) help to identify future research avenues.

Our findings underscore the importance of identifying the salient characteristics of the nighttime visual landscape for pumas. Choosing darker parts of the landscape to cross roads suggests that there are aspects of the lightscape that pumas attend to. It is possible that pumas are responding to local light conditions but not to average light conditions as measured here; there are other characteristics of light that may be responsible for driving puma behavior that were not specifically the subject of this study. In addition to average luminance above and below the road verge, other aspects of the lightscape might include 1) variation between the brightest and the darkest parts of the lightscape (i.e., contrast), 2) distribution of lights across the lightscape, 3) light spectra weighted for puma vision since luminance is a human-biased metric, and 4) landscape levels of radiance (light from a source) versus irradiance (light incident on a surface). For example, light arrangement on the landscape can lead to high contrast where a well-lit road can make it difficult to discern the landscape in adjacent dark areas (Rich & Longcore 2013).

Identifying optimal measurements of electromagnetic radiation relevant to wildlife behavior is a challenge (Johnsen 2012). There are two main areas of complexities that future research might address: specific metrics used in quantifying the lightscape and assessing salient aspects of the lightscape important for shaping wildlife behavior. We present here human generated night-time light levels across the sky from a puma eye-level field of view; behavior could also be driven by other metrics of the lightscape including, for example, overhead skyglow (e.g., influencing the ability to see, and be seen by, prey), discrete bright lights on the horizon (e.g., affecting orientation or navigation), contrast (e.g., bright road lights inhibit the ability to see through a dark wildlife undercrossing), or light variation in the scene (e.g., the role of light from moving vehicle headlights). Additionally, response to any of these aspects of light in the view-shed may shift based on behavioral state (Zeller et al. 2014). Possible approaches for addressing these questions could include coupling animal-borne light measuring equipment with roadside behavior (derived from high temporal resolution GPS collars). Road crossing locations, and locations where animals approach the road but did not cross (again from high temporal resolution GPS collar data), could later be assessed for a host of lightscape variables.

Our finding that light, from a puma eye-level perspective, did not influence puma crossing intensity could point to several underlying mechanisms. The first is that there is an underlying correlation between puma crossing intensity and light, however, there was insufficient data to find statistical significance. A second possibility is that pumas were not using light at this very local scale to make decisions on where to cross secondary roads, or that other factors were more important such as a strong motivation to cross that would overwhelm potential light effects. A third possibility is that a single value, mean radiance across the field-of-view, is insufficient to describe the lightscape above and below the road verge. Finally, a fourth possibility is that light at the roadside must be measured at different distances from the roadside to better reflect what puma see as they approach the road rather than the lightscape at the immediate roadside. Future research could focus on measuring light across a diversity of road sizes, quantifying multiple metrics of light at the roadside (e.g., hemispherical photography, sky glow, scene contrast, temporal variability, presence/absence of bright lights on the horizon, road illumination), and measuring light along the path that a puma experiences before approaching a road. Including light meters on GPS collars could be a productive methodology for future studies focused on light at the roadside.

### 4.1 Conservation impact

Vehicular collisions are an important source of mortality for some species, including the pumas that we studied here. Reducing wildlife-vehicle collisions and increasing use of wildlife crossing structures requires an understanding of stimuli that both attract and repel individuals. Transportation agencies have few tools and guiding principles for identifying optimal locations for expensive wildlife crossing locations – this research provides insight on examining landscapes for locating mitigation structures.

We have shown that pumas cross roads in areas with relatively less anthropogenic light. This has concrete implications for identifying optimal future crossing structure locations.

Knowledge of puma unwillingness to cross sections of highway with particular light profiles, from results presented here coupled with future research, can be used in two ways. Wildlife crossing structures, designed to permit safe passage across a road, can be located and designed using “puma-friendly” lighting to encourage use. Alternately, lighting profiles that pumas avoid could be used to repel pumas from road stretches that are hotspots for puma-vehicle collisions. Identifying precisely how anthropogenic stimuli both attract and repel animals (Greggor et al. 2020) creates useful tools to potentially modify their movement.

## Impact statement

Human generated light at night may influence puma movement behavior and influences where puma cross roads.

## Acknowledgments

H.B. Shaffer, A. Wolf, M. Colon, P. Hiscocks, T. Longcore, J. Valler, K. Zeller, L. Lee, D. Kamradt, B. Clarke, R. Klufas, J. Milligan, C. Kyba, Y. Yang, M. Yang, T. Song, Yang Yang Lab, T. Saggese, S. Xu, Southern California Edison.

## Supporting Information

Data file with field site locations, descriptive information, and site-specific data collected (Appendix S1) is available online. The authors are solely responsible for the content and functionality of these materials. Queries (other than absence of the material) should be directed to the corresponding author.

## Data Availability Statement

Data used in this manuscript are available in the supplemental information (Appendix S1)

## Author contributions

EA, SR, and DB conceived the ideas and designed methodology; EA, JS, and SR collected the data; EA analysed the data; EA led the writing of the manuscript. All authors contributed critically to the drafts and gave final approval for publication.

## Literature Cited

Benson JF, Sikich JA, Riley SPD. 2020. Survival and competing mortality risks of mountain lions in a major metropolitan area. Biological Conservation 241:108294.

car package. 2020. John Fox and Sanford Weisberg (2019). An {R} Companion to Applied Regression, Third Edition. Thousand Oaks CA: Sage. URL: https://socialsciences.mcmaster.ca/jfox/Books/Companion/.

Conover WJ. 1998. Practical nonparametric statistics. John Wiley & Sons.

Elvidge CD, Baugh K, Zhizhin M, Hsu FC, Ghosh T. 2017. VIIRS night-time lights. International Journal of Remote Sensing 38:5860–5879.

Forman RT, Sperling D, Bissonette JA, Clevenger AP, Cutshall CD, Dale VH, Fahrig L, France RL, Goldman CR, Heanue K. 2003. Road ecology: science and solutions. Island press.

GDT Dynamap. 2014. SteetMap07 Roads. Tele Atlas North America, Inc. Available from http://www.teleatlas.com.

Greggor AL, Berger-Tal O, Blumstein DT. 2020. The Rules of Attraction: The Necessary Role of Animal Cognition in Explaining Conservation Failures and Successes. Annual Review of Ecology, Evolution, and Systematics 51:483–503.

gvlma package. 2019. Edsel A. Pena and Elizabeth H. Slate. gvlma: Global Validation of Linear Models Assumptions. R package version 1.0.0.3. https://CRAN.R-project.org/package=gvlma.

Hiscocks PD. 2011, January. Measuring Light. Syscomp Electronic Design Limited, Toronto. Available from http://www.ee.ryerson.ca/~phiscock/.

Hiscocks PD. 2013a, October 30. Integrating Sphere for Luminance Calibration. Syscomp Electronic Design Limited, Toronto. Available from http://www.ee.ryerson.ca/~phiscock/astronomy/light-pollution/integrating-sphere.pdf (accessed February 10, 2015).

Hiscocks PD. 2013b, November 25. Measuring Luminance with a Digital Camera: Case History. Syscomp Electronic Design Limited, Toronto. Available from http://www.ee.ryerson.ca/~phiscock/astronomy/light-pollution/luminance-case-history.pdf (accessed January 5, 2015).

Hiscocks PD. 2014, February 16. Measuring Luminance with a digital camera. Syscomp Electronic Design Limited, Toronto. Available from www.syscompdesign.com (accessed February 10, 2015).

Hu Z, Hu H, Huang Y. 2018. Association between nighttime artificial light pollution and sea turtle nest density along Florida coast: A geospatial study using VIIRS remote sensing data. Environmental Pollution 239:30–42.

Johnsen S. 2012. The Optics of Life: A Biologist’s Guide to Light in Nature. Princeton University Press.

Lazo-Cancino D, Musleh SS, Hernandez CE, Palma E, Rodriguez-Serrano E. 2017. Does silvoagropecuary landscape fragmentation affect the genetic diversity of the sigmodontine rodent Oligoryzomys longicaudatus? PeerJ 5:e3842. PeerJ Inc.

Levin N et al. 2020. Remote sensing of night lights: A review and an outlook for the future. Remote Sensing of Environment 237:111443.

National Park Service. 2014. Geospatial data for the Vegetation Mapping Inventory Project of Santa Monica Mountains National Recreation Area. Santa Monica Mountains and Environs, California, USA.

Ng SJ, Dole JW, Sauvajot RM, Riley SPD, Valone TJ. 2004. Use of highway undercrossings by wildlife in southern California. Biological Conservation 115:499–507.

R Core Team. 2020. R 4.0.2: a language and environment for statistical computing. R Foundation for Statistical Computing, Vienna, Austria. Available from http://www.R-project.org.

Rasband W. (n.d.). ImageJ. National Institutes of Health, USA. Available from https://imagej.nih.gov/.

Reed DF, Woodard TN. 1981. Effectiveness of Highway Lighting in Reducing Deer-Vehicle Accidents. The Journal of Wildlife Management 45:721–726.

rgeos package. 2020. Roger Bivand and Colin Rundel (2020). rgeos: Interface to Geometry Engine - Open Source (‘GEOS’). R package version 0.5-5. https://CRAN.R-project.org/package=rgeos.

Rich C, Longcore T. 2013. Ecological Consequences of Artificial Night Lighting. Island Press.

Riley SPD, Brown JL, Sikich JA, Schoonmaker CM, Boydston EE. 2014a. Wildlife Friendly Roads: The Impacts of Roads on Wildlife in Urban Areas and Potential Remedies. Pages 323–360 in R. A. McCleery, C. E. Moorman, and M. N. Peterson, editors. Urban Wildlife conservation. Springer US. Available from http://link.springer.com/chapter/10.1007/978-1-4899-7500-3_15 (accessed April 17, 2015).

Riley SPD, Pollinger JP, Sauvajot RM, York EC, Bromley C, Fuller TK, Wayne RK. 2006. A southern California freeway is a physical and social barrier to gene flow in carnivores. Molecular Ecology 15:1733–1741.

Riley SPD, Sauvajot RM, Fuller TK, York EC, Kamradt DA, Bromley C, Wayne RK. 2003. Effects of Urbanization and Habitat Fragmentation on Bobcats and Coyotes in Southern California. Conservation Biology 17:566–576.

Riley SPD, Serieys LEK, Pollinger JP, Sikich JA, Dalbeck L, Wayne RK, Ernest HB. 2014b. Individual Behaviors Dominate the Dynamics of an Urban Mountain Lion Population Isolated by Roads. Current Biology 24:1989–1994.

Robertson BA, Hutto RL. 2006. A framework for understanding ecological traps and an evaluation of existing evidence. Ecology 87:1075–1085.

Russell R. 2016. Canon Hack Development Kit (CHDK). Available from http://www.rupert.id.au/tutorials/CHDK/index.php.

Santa Monica Mountain Recreation Area vegetation map, super-generalized version. 2015. Based on SAMO’s 2007 vegetation map, the super-generalized version uses generalized vegetation classes developed by Brendan Clarke and Robert S Taylor, NPS-SAMO. Available from https://apps.wildlife.ca.gov/bios.

SCAG. 2005. Land-use data for Ventura and Los Angeles counties. Southern California Association of Goverments, Los Angeles, California.

Smith JA, Wang Y, Wilmers CC. 2015. Top carnivores increase their kill rates on prey as a response to human-induced fear. Proceedings of the Royal Society of London B: Biological Sciences 282:20142711.

Soulé ME. 1985. What Is Conservation Biology? BioScience 35:727–734.

USGS. 2015. Digital Elevation Model, region 4. United States Geological Survey. Available from https://viewer.nationalmap.gov/basic/#/.

Vickers TW, Sanchez JN, Johnson CK, Morrison SA, Botta R, Smith T, Cohen BS, Huber PR, Ernest HB, Boyce WM. 2015. Survival and Mortality of Pumas (Puma concolor) in a Fragmented, Urbanizing Landscape. PLOS ONE 10:e0131490.

Wade AA, McKelvey KS, Schwartz MK. 2015. Resistance-surface-based wildlife conservation connectivity modeling: Summary of efforts in the United States and guide for practitioners. Available from http://www.treesearch.fs.fed.us/pubs/48464 (accessed January 14, 2016).

Witherington BE. 1997. The problem of photopollution for sea turtles and other nocturnal animals. Pages 303–328 Behavioral approaches to conservation in the wild. Cambridge University Press Cambridge.

Zeller KA, McGarigal K, Beier P, Cushman SA, Vickers TW, Boyce WM. 2014. Sensitivity of landscape resistance estimates based on point selection functions to scale and behavioral state: pumas as a case study. Landscape Ecology 29:541–557.

Zollner PA, Lima SL. 1999. Illumination and the perception of remote habitat patches by white-footed mice. Animal Behaviour 58:489–500.

